# Developmental Reprogramming of Hypothalamic-Pituitary Axis in Mice by Common Environmental Pollutants

**DOI:** 10.1101/2024.01.04.574224

**Authors:** Joshua P. Mogus, Marjorie Marin, Olatunbosun Arowolo, Victoria Salemme, Alexander Suvorov

**Author notes:** Corresponding author and person to whom reprint requests should be addressed: Alexander Suvorov, School of Public Health & Health Sciences, Department of Environmental Health Sciences, University of Massachusetts – Amherst, 240B Goessmann, 686 N. Pleasant Street, Amherst, MA 01003 USA, **Email:**.

## Abstract

Humans are exposed to a large number of endocrine disrupting chemicals (EDCs). Many studies demonstrated that exposures to EDCs during critical windows of development can permanently affect endocrine health outcomes. Most of experimental studies address changes in secretion of hormones produced by gonads, thyroid gland and adrenals, and little is known about the ability of EDCs to produce long-term changes in the hypothalamic-pituitary (HP) control axes. Here, we examined the long-term effects of three common EDCs on male mouse HP gene expression, following developmental exposures. Pregnant mice were exposed to 0.2 mg/ml solutions of bisphenol S (BPS), 2,2’,4,4’-tetrabromodiphenyl ether (BDE-47), or 3,3’,5,5’-tetrabromobisphenol A (TBBPA) from pregnancy day 8 through lactation day 21 (weaning day). Male offspring were left untreated until postnatal day 140, where pituitaries and hypothalami were collected. Pituitaries were assed for gene expression via RNA sequencing, while specific genes were assessed for expression in hypothalami via RT-qPCR. Differential expression, as well as gene enrichment and pathway analysis, indicated that all three chemicals induced long-term changes, (mostly suppression) in pituitary genes involved in its endocrine function. BPS and BDE-47 produced effects overlapping significantly at the level of effected genes and pathways. All three chemicals altered genes and pathways of gonad and liver HP axes, while BPS altered HP-adrenal and BDE-47 altered HP-thyroid pathways specifically. All three chemicals also reduced expression of immune genes in the pituitaries. Targeted gene expression in the hypothalamus indicates a down regulation of hypothalamic endocrine control genes by BPS and BDE-47 groups, concordant with changes in the pituitary and suggests that these chemicals suppress the overall HP endocrine function. Interestingly, all three chemicals altered pituitary genes of GPCR-mediated intracellular signaling molecules, many of which are key signalers common to many pituitary responses to hormones. The results of this study show that developmental exposures to common and ubiquitous EDCs have long-term impacts on hormonal feedback control at the hypothalamic-pituitary level.

## Introduction

Today, humans are exposed to a broad variety of environmental pollutants ^1–3^, many of which have been identified as endocrine-disrupting chemicals (EDCs), or chemicals that alter one or more aspects of hormone signaling ^4^. The early development of healthy and functional hypothalamic-pituitary axes (HP axes) is critical to the establishing and maintaining endocrine homeostasis in adulthood ^5^. Nearly all organ systems have some feedback regulation by HPs including the gonads, thyroid, adrenals, and liver ^6^. However, exposure to EDCs during critical developmental windows can alter the organization and regulation of hormone systems permanently, leading to greater risks for later disease ^7^. The majority of EDC studies using *in vivo* models typically evaluate endocrine function at lower levels of the HP axis hierarchy, (i.e. organs and downstream tissues) ^8,9^. Though important, these studies cannot discern if changes in the organ/tissue are from direct activity of the chemical in the target tissue, or if the tissue is responding to a disruption of upstream HP control. Ultimately, there is still a paucity of knowledge on how the pituitary and hypothalamus respond to developmental EDC exposures and how these changes may affect HP feedback functions.

Here we evaluate the long-term effects of developmental exposure to either bisphenol-S (BPS), 2,2’,4,4’- tetrabromodiphenyl ether (BDE-47), or 3,3’,5,5’-tetrabromobisphenol A (TBBPA), three common EDCs, on brain HP axis gene expression in the CD-1 mouse model. BPS is a sulfonated bisphenol and replacement of the well-recognized EDC and plastic component, bisphenol-A (BPA). Similar to BPA, BPS demonstrates estrogenic effects ^10^ and has been shown to disrupt several endocrine endpoints of development including metabolism ^11^, mammary gland structure ^12,13^, and behavior ^14,15^. BDE-47 is a polybrominated diphenyl ether (PBDE), used as flame retardant additive in a variety of polymers. Although, commercial use of PBDE has ceased in many developed countries, their concentrations remain high in human samples in the North America ^16,17^. Developmental exposures to BDE-47 in laboratory animals and humans result in the disruption of lipid and carbohydrate metabolism ^18–22^, and deterioration of reproductive health ^23^ and impairment of cognitive development ^24,25^. PBDEs are known also to disrupt multiple endocrine mechanisms, including the direct interaction with hormonal receptors, transporting proteins, and the induction of metabolic enzymes responsible for the metabolism of hormones ^26^. For example, PBDEs disrupt hormonal signaling through thyroid and steroid axes ^27^ and interfere with the insulin-like growth factor (IGF) axis ^19,28^. TBBPA is a replacement brominated flame retardant, which has been shown to alter thyroid axis signaling ^29^, steroidogenesis ^30^, spermatogenesis ^31^ and affect endocrine mediated behaviors ^15,32,33^.

In the current study, we demonstrate long-term changes in gene expression in hypothalamus and pituitary in mice perinatally exposed to environmentally relevant doses of all tested compounds, indicative of altered hormonal control by HP axes. These findings suggest that dysregulation of the HP control may be a common effect of EDCs.

## Methods

### Animals and Treatment

Eight-week-old male and female CD-1 mice were purchased from Charles River Laboratories (Kingston, NY, USA) and housed at the University of Massachusetts - Amherst animal care facility. Mice were housed in polysulfone cages with controlled temperature (23 ±2 °C), humidity (RH% 40 ± 10) and lighting (12h dark light cycle). The animals were provided fresh water and mouse chow (LabDiet Chow 5058; LabDiet, St. Louis, MO) *ad libitum*. Males and females were placed in separate cages with no more than four individuals per cage. Males and females were left to acclimate separately for three days before being set together for mating. Daily checks for vaginal plugs were conducted, with the date of plug detection indicating pregnancy day one (P1). Pregnant females were placed into one of the four following treatment groups: tocopherol stripped corn oil (Miomedicals, Solon, OH), 0.2 mg/ml solution of TBBPA Sigma-Aldrich, 97% purity) in corn oil, 0.2 mg/ml solution of BDE-47 (Accustandard, Inc., New Haven, 100% purity) in corn oil, or 0.2 mg/ml solution of BPS (Santa Cruz Biotechnology, Dallas, TX, 99% purity) in corn oil.

Starting from pregnancy day eight (P8) until lactation day 21 (LD21) F0 females were treated with their respective solutions once a day, every day (Figure 1). Before each treatment, females were weighed, and solutions were calculated to provide 1 µl/gram body weight (BW) to equal 0.2 mg/kg BW/day exposure for the respective treatment. The solutions were delivered by pipette feeding ^34^. Dams were left to deliver offspring and nurse naturally. No pups were culled to maintain consistency of nutrient distribution among the same number of fetuses/pups at pre- and postnatal periods and avoid catch-up growth ^35^. On PND21, pups were removed from their mothers, separated by sex, and then placed in new cages with no more than 4 individuals per cage. Over the course of ten weeks, starting PND70, male offspring were tested for social behavior. The results of behavioral testing and analysis of fecal microbiome changes in response to exposures were previously published ^15,36^. All male animals were euthanized on PND140 via decapitation. Hypothalami and pituitaries were immediately collected upon euthanasia, snap-frozen with liquid nitrogen, and stored at -80°C. Female offspring were not included in this study as in females expression of endocrine genes may be effected by estrus cycle. All care and experimental procedures involving animals met the requirements of the National Institute of Health’s Guide for Care and Use of Laboratory and were approved by the Institutional Animal Care and Use Committee at the University of Massachusetts – Amherst (Protocol # 2013-0069).

**Figure 1:**
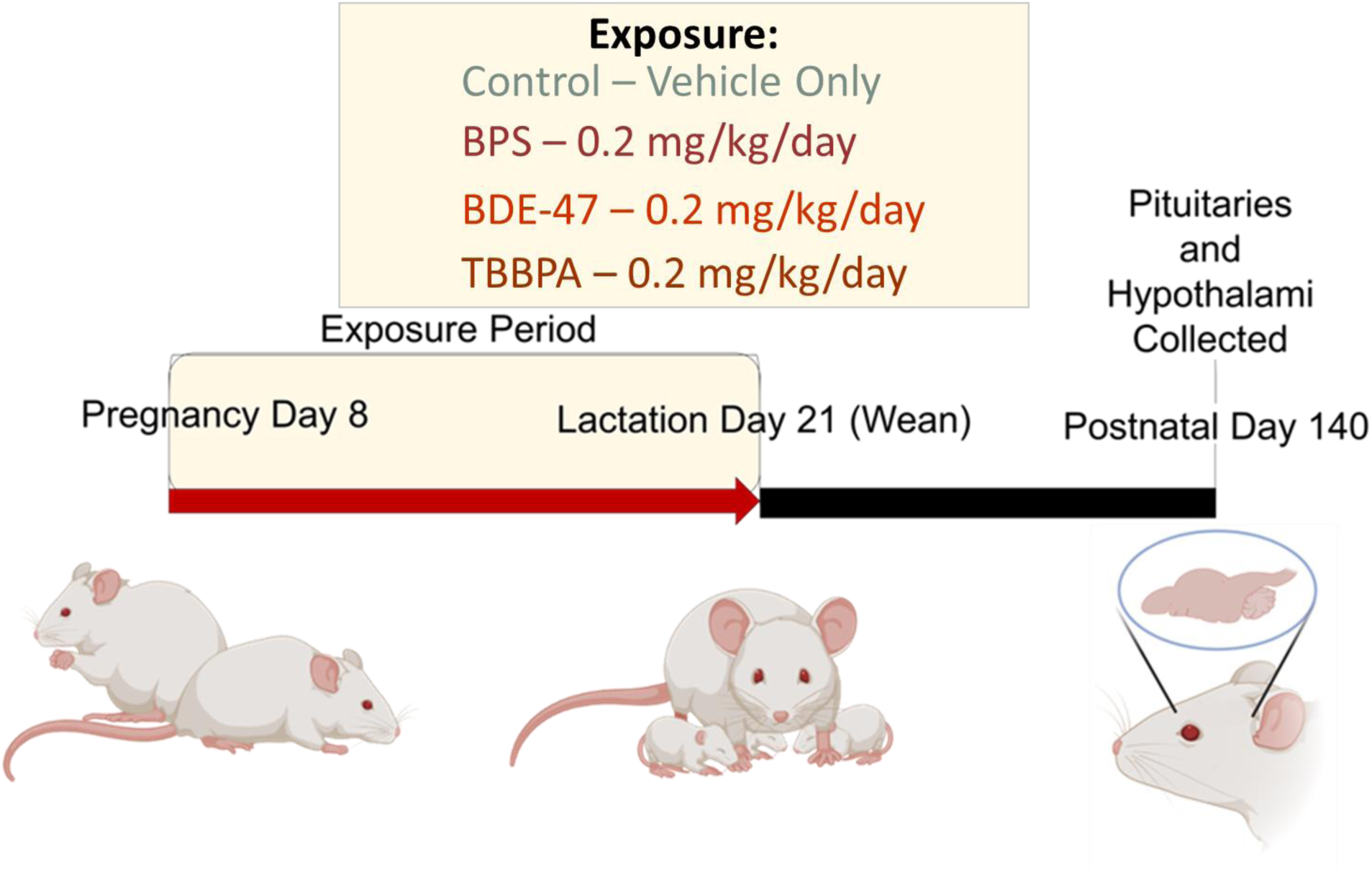
Schematic of experimental design.

### RNA extraction and Isolation

RNA was collected using a standard Trizol extraction procedure according to manufacturer’s guidelines. Whole pituitaries or hypothalami were homogenized in Trizol (Ambion, Life technologies or Invitrogen) using a handheld rotor homogenizer over ice. Homogenates were phase separated via chloroform, precipitated with isopropanol, and washed two times with 75% ethanol. RNA precipitate was resuspended into RNase-free Tris-EDTA (TE) buffer. All samples underwent quality control checks; Nanodrop 1000 (Thermo Fisher Scientific, Wilmington, DE) for RNA concentration and purity and Agilent 2100 Bioanalyser (Agilent Technologies, Santa Clara, CA) for integrity via 28s ribosomal subunit-based RNA Integrity Number (RIN) analysis. Only samples with RIN values >9 were used for library preparation.

### Library Prep for RNAseq

To evaluate global mRNA expression within the pituitary, we used the standard Illumina TruSeq protocol (TruSeq Stranded mRNA LP, Cat # 20020594 and TruSeq RNA Sg Idx SetB, Cat # 20020493, Illumina, San Diego, CA)) following the manufacturers recommendations and as was done previously ^31^. Briefly, intact mRNA with polyadenylated tails were isolated from 1 µg mRNA samples. Intact mRNA was then used to generate strand-specific libraries with multiplexing indexes. cDNA libraries were built to fragments lengths of 300 base pairs. Quality and purity of libraries were evaluated using the Agilent 2100 Bioanalyzer and library concentrations and purities measured using a Qubit 3.0 fluorometer (Life Technologies, Carlsbad, CA). High-throughput sequencing was conducted using the NextSeq500 sequencing system (Illumina, San Diego, CA) in the genomic core facility of the University of Massachusetts Amherst. cDNA libraries were singe-end sequenced in 76 cycles using a NSQ 500/550 Hi Output KT v2.5 (Cat #20024906 Illumina, San-Diego, CA) in one multiplex run (N=3/exposure group). Sequencing data are available via Zenodo public repository ^37^.

### RT-qPCR

RT-qPCR was used to validate RNA-seq results for pituitary and to analyze expression of select genes in hypothalamus. Total RNA was purified of genomic DNA contamination using DNase (RQ1 RNAse-free DNAse, Cat. # M610A, Promega, Madison, WI) and reverse transcribed using the High Capacity cDNA Reverse Transcription Kit (Cat.# 4368814, Applied Biosystems, Vilnus, Lithuania). Using free online software Primer3Plus (http://primer3plus.com/cgi-bin/dev/primer3plus.cgi), forward and reverse primers were designed to anneal different exons spanning at least one intron. Triplicate 5-μl real-time PCR reactions, each containing iTaq Universal SYBR Green Supermix (Cat 172–5124, BioRad), primers, and cDNA template were loaded onto a 384-well plate and run through 40 cycles on a CFX384 real time cycler (Bio-Rad Laboratories, Inc). The data were analyzed using the manufacturer’s CFX manager software, version 3.1. Relative quantification was determined using the ΔΔCq method (Livak and Schmittgen, 2001). We selected two housekeeping genes (HKGs), *B2M* [beta-2-mircoglobulin] and *Tbp* [TATA box-binding protein like 1], due to their validated expression stability across body tissues, and accepted use a HKGs in a wide body of literature ^38^. ΔΔCt was normalized to the Tukey’s biweight centered means of the HKG values. *B2M* and *Tbp* were validated for stable expression using RNA sequencing in pituitary, where in all cases, treatment did not significantly alter HKG expression. Primer sequences for qPCR provided in (*Supplemental Table 1*).

### Analysis of RNA Sequencing

Read filtering, trimming, and de-multiplexing were performed using the BaseSpace cloud service by Illumina (https://basespace.illumina.com/home/index, RRID:SCR_011881). Processed reads were mapped to the mouse reference genome (MM10) using the RNA-Seq Alignment v. 1.1.1. software with Bowtie 2 and aligned reads were used to assemble transcripts and analyze differential expression using Cufflinks Assembly & DE v. 2.1.0. package. The short-lists of differentially expressed genes (DEGs) (FDR q ≤ 0.05) were used for the analysis of enriched biological categories using Metascape and Ingenuity Pathway Analysis with default settings ^39^. IPA Upstream Regulators was conducted to predict likely activity changes to transcription regulators upstream of our observed DEGs. Here, IPA calculates a z-score to predict the activational state of the upstream regulator as either “activated” or “inhibited” based on observed differential gene expression in our treated samples. Activational z-scores greater than 2.0 or smaller than -2.0 are considered statistically significant (p < 0.05). Metascape analysis was conducted separately for positively and negatively regulated DEGs. Files for differential expression, IPA canonical pathways, and Metascape gene enrichment analysis are available via Zenodo ^37^.

## Results

There was no significant difference in mean litter sizes between exposure groups. No weight differences were observed for control and exposed dams throughout pregnancy and lactation. There were also no significant changes in average pup weight adjusted for litter size at birth [this information was previously published ^15^].

Sequencing was completed with an average of 35 million reads per sample. The percentage of reads that aligned to the reference genome varied between 80.5 and 95.5 across samples, with percent stranded reads around 98 for all samples. Differential expression values were identified for 16,393, 16,300 and 15,935 unique assembled transcripts with known identifiers (genes, pseudogenes, non-coding RNA) in TBBPA, BPS and BDE-47 exposure groups respectively. RNA-seq results were validated by RT-qPCR for select genes, *Krt17* [Keratin, type I cytoskeletal 17] and *Vgf* [Vascular endothelial growth factor A] for BPS and BDE-47 exposures (*Supplemental File 1*).

### BPS induces long-term changes in HP hormone signaling pathways

48 differentially expressed genes (DEGs) within the pituitary were identified with FDR q ≤ 0.05 *Supplemental File 2,* also in repository ^37^). While 42 of these DEGs were downregulated, only six were upregulated. Several of the downregulated DEGs are known to be associated with classic pituitary endocrine function: *Egr1* [Early growth response protein 1], the zinc-finger transcription figure that regulates gonadotropin signaling ^40^, *Lhb* [Luteinizing hormone beta] which codes for the beta subunit of the gonad-stimulating luteinizing hormone ^41^, *Gh* [Growth hormone/Somatostatin] which transcribes growth hormone, a key hormonal regulator of growth and metabolism ^42^, and *Nr4a1* [Nuclear receptor subfamily 4 group A member 1], a pituitary mediator of stress known to be responsive to corticotropin releasing hormone (CRH), and a transcription factor that regulates differentiation and apoptosis in macrophages and T cells. ^43–45^. Additionally, *Il36G* [Interleukin-36 gamma], an important molecule in immune response signaling, was also downregulated. Notably, it was observed a downregulation of intracellular signaling molecules that have phosphorylation or transcription factor functions associated with growth factor and G-protein coupled receptor (GPCR) activity: *Rasd1* [Dexamethasone-induced Ras-related protein 1], *Dusp1* [Dual specificity protein phosphatase 1]*, Fos* [Protein c-Fos], and *JunB* [Transcription factor Jun B].

To further evaluate differentially regulated functions in the pituitary, Metascape pathway enrichment analysis was conducted with our short list of 48 DEGs (*Supplemental File* 3, also in repository ^37^). Several biological categories relevant to endocrine function were negatively enriched by BPS including, response to hormone, negative regulation of stress-activated MAPK cascade, positive regulation of cell death, intracellular signaling pathway, and post-embryonic development (*Figure 2A*). Biological categories with a positive enrichment with genes activated by BPS include retina vasculature development in camera-type eye, and blood vessel development (*Figure 2B*).

**Figure 2:**
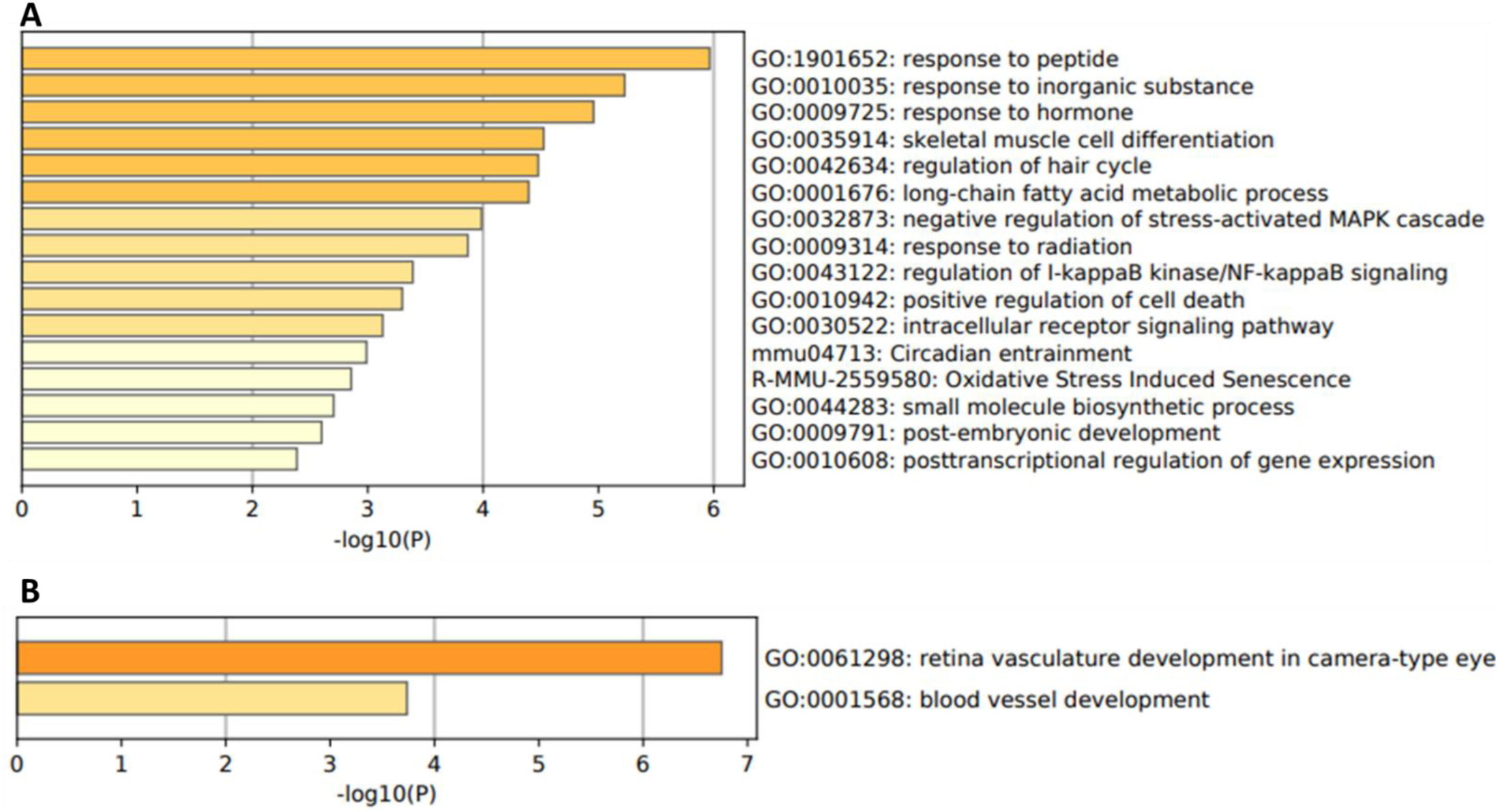
BPS gene ontology analysis. Top Metascape biological categories that were (**A**) negatively enriched by BPS exposure or (**B**) positively enriched.

IPA analysis identified 79 canonical pathways significantly (-log p-values of ≥1.3) enriched with pituitary DEGs (*Supplemental File 4*, also in repository ^37^). All enriched pathways were predicted to be negatively regulated by BPS exposure. BPS enriched several pathways associated with pituitary endocrine function (*Table 1*), including important hormonal signaling pathways: GNRH signaling, glucocorticoid receptor signaling, and growth hormone signaling among others. Of note, several altered pathways were directly associated with immune function (at least 28), while 71 out of 79 altered pathways included at least one generalized intracellular phosphorylation cascade molecule (*Fos*, *Rasd1,* or *Junb*).

**Table 1:** Endocrine function-related IPA pathways enriched in pituitary by BPS exposure. IPA canonical pathway analysis of differentially expressed genes from BPS exposed mice to controls. Significantly enriched pathways determined by log p-values ≥ 1.3. All pathways were predicted to be downregulated.

| Ingenuity Canonical Pathway | P-value | Differentially Expressed Genes |
| --- | --- | --- |
| GNRH Signaling | 0.0002 | *Egr1, Fos, Lhb, Rasd1* |
| PPAR Signaling | 0.0006 | *Fos, Il36g, Rasd1* |
| Glucocorticoid Receptor Signaling | 0.0021 | *Dusp1, Fos, Krt17, Rasd1* |
| ERK/MAPK Signaling | 0.0038 | *Dusp1, Fos, Rasd1* |
| Growth Hormone Signaling | 0.0060 | *Cshl1, Fos, Gh* |
| Corticotropin releasing hormone signaling pathway | 0.0240 | *Fos, Nr4a1* |
| Prolactin Signaling | 0.0080 | *Fos, Rasd1* |
| IGF Signaling | 0.0126 | *Fos, Rasd1* |
| Thyroid Cancer Signaling | 0.0074 | *Fos, Rasd1* |
| Estrogen Dependent Cancer Signaling | 0.0065 | *Fos, Rasd1* |
| Ovarian Cancer Signaling | 0.0214 | *Lhb, Rasd1* |

IPA Upstream analysis was performed to evaluate predicted upstream regulators that may explain our altered DEGs and pathways. We conducted an upstream analysis against endogenous and pharmaceutical regulators (*Table 2*). All identified upstream regulators were predicted to be inhibited, including several regulators associated with HP axis hormone signaling. Most of these were directly associated with estrogen signaling. Indeed, beta-estradiol (aka estradiol or 17beta-estradiol), is an endogenous hormone and the primary ovarian estrogen and agonist of ER and membrane ER (mER). 4-hydroxytamoxifen is an active metabolite of the pharmaceutical estrogen tamoxifen and a known selective estrogen receptor modifier. GPER1 (G protein-coupled estrogen receptor 1), an estrogen receptor responsible for the rapid non-genomic response to estrogen, was predicted to be inhibited. In connection to this, IPA predicted a reduction in forskolin activity, a pharmaceutical chemical known to increase cAMP activity. Additionally upstream regulator analysis predicted inhibition of GnRH receptor agonist (GnRH-A) activity and activity of a generalist transcription factor, CREB1.

**Table 2:** Top predicted IPA upstream regulators for BPS exposed pituitaries.

| Upstream Regulator | Predicted Effect | Z-Score Activational Value |
| --- | --- | --- |
| Forskolin | Inhibited | -4.169 |
| CREB1 | Inhibited | -4.022 |
| 4-hydroxytamoxifen | Inhibited | -3.063 |
| GnRH-A | Inhibited | -2.941 |
| GPER1 | Inhibited | -2.733 |
| Beta-estradiol | Inhibited | -2.413 |

To assess if changes in hypothalamus hormone signaling pathways were altered in conjunction with pituitary hormone pathways, we used targeted RT-qPCR of selected hormone signaling genes (*Figure 3*). Following the IPA suggested altered downstream pathway analysis, we first looked at expression of gonadotropin and HP-gonad axis pathway molecules. We evaluated gonadal hormone receptors that provide the principal recognition of circulating steroids in the hypothalamus. Of these we found that nuclear estrogen receptor, *Esr1* [Estrogen receptor alpha] was significantly downregulated due to BPS exposure, while the estrogen membrane receptor (*Gper1*), and nuclear progesterone receptor beta (*Pgrβ)* were not changed.

**Figure 3:**
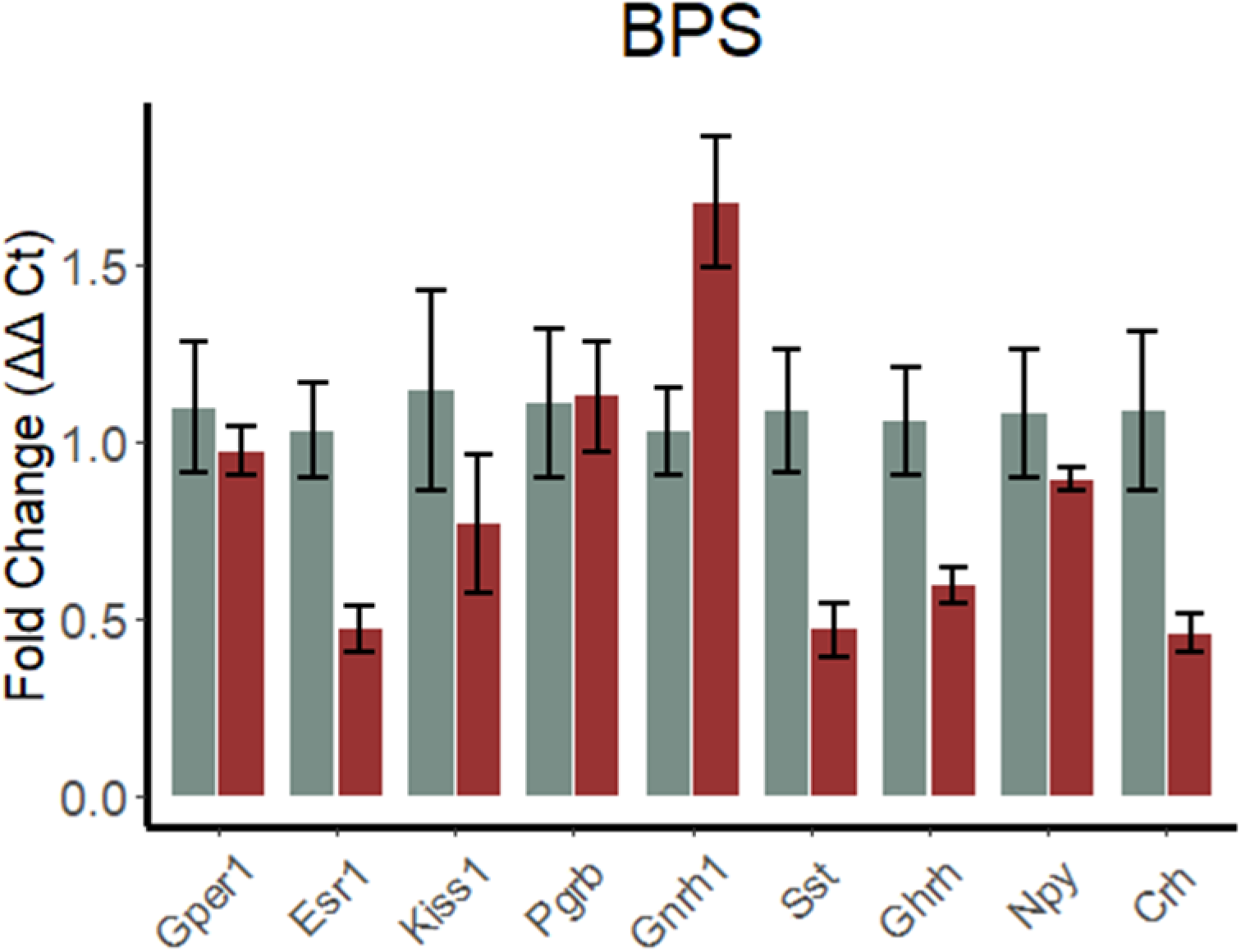
BPS alters long-term hormone gene expression in the hypothalamus. Gene expression of key hormone regulating genes in hypothalamus via RT-qPCR. Gray bars signify controls, while maroon are for BPS group. Significance determined by Student’s t-test or Welch’s -test where relevant, α = 0.05, * represents p < 0.05. Error bars represent standard error.

Next, we evaluated gene expression of *Kiss1* [Metastasis-suppressor KiSS-1], an important intermediary molecule between hypothalamic steroid receptors and GnRH. Though differences in *Kiss1* expression were not significant, the downstream *Gnrh1* [Progonadoliberin-1/Gonadotropin-releasing hormone) was significantly upregulated by BPS.

To assess if altered pathways in HP-Liver signaling were matched in the hypothalamus, we measured gene expression of the somatotropic regulator, *Ghrh* [Somatoliberin/Growth hormone-releasing hormone] and its major inhibitor *Sst* [Somatostatin]. Interestingly, both molecules were significantly downregulated by BPS exposure. Last, we assessed hypothalamic regulators of pituitary glucocorticoid function. Here, we found that *Crh* [Corticoliberin/Corticosteroid -releasing hormone], the critical hormone that induces corticotropic hormones in the pituitary, was significantly downregulated. Conversely, its upstream activator in the hypothalamus, *Npy* [Pro-neuropeptide Y], was not altered.

### BDE-47 induces long-term changes in HP hormone signaling pathways

317 DEGs were identified in BDE-47 exposed mouse pituitaries using our selection criteria of q value ≤ 0.05 (*Supplemental File 5*, also in repository^37^). Of the 317 DEGs, 182 DEGs were significantly downregulated, while 135 were upregulated. Several affected genes were a part of pituitary endocrine function. These DEGs included: down-regulation of *Egr1*, *Tshβ* [Thyrotropin subunit beta], and *Dio1* [Type I iodothyronine deiodinase]. Additionally, there was an increase in expression of *Gnrhr* [gonadotropin-releasing hormone receptor]. Like the BPS group, there were DEGs known to participate in immune response such as the gene encoding the interleukin family receptor *iL6R* [Interleukin-6 Receptor], and the interleukin, *Il36G.* Generalized intracellular phosphorylating signaling molecules *Fos*, *JunB*, *Rasd1*, *Gab1* [Growth factor receptor bound protein 2 associated protein 1], and *Dusp1* were also downregulated. Furthermore, genes for paracrine (secreted protein) signaling pathways were downregulated. These included the TGF-β [Transforming growth factor beta] signaling molecules, *Bmp7* [Bone morphogenic protein 7] and *Vdr* [Vitamin D receptor], as well as Wnt [Wingless]/β-catenin signaling molecules *Sfrp5* [Secreted frizzled-related protein 5], *Sox13* [SRY-Box transcription factor 13], *Sox4* [SRY-Box transcription factor 4], *Fzd1* [Frizzled class receptor 1], *Wnt10A* [Wnt Family member 10A], and *Ubc* [Ubiquitin C].

To further evaluate biological functions differentially regulated by BDE-47 exposure, Metascape pathway enrichment analysis was conducted with the our short list of 317 DEGs differentially expressed genes in the pituitary (full list in *Supplemental File 6*, also in repository ^37^). The top biological pathways negatively enriched by BDE-47 exposures are shown in (*Figure 4A*). Some enriched categories were relevant for the endocrine function of pituitary, including *Negative regulation of phosphorylation, Epithelial cell differentiation, Response to corticotropin-releasing hormone, Regulation of hormone levels, and Regulation of ERK1 and ERK2 cascade*. See ^37^ for the full list of negatively enriched categories. In addition, 14 positively enriched pathways were associated with BDE-47 exposure (*Figure 4B*) with some having relation to pituitary endocrine function: *Regulation of epithelial cell differentiation, Regulation of peptide transport, HSP90 chaperone cycle for steroid hormone receptors (SHR), and Endocrine pancreas development*.

**Figure 4:**
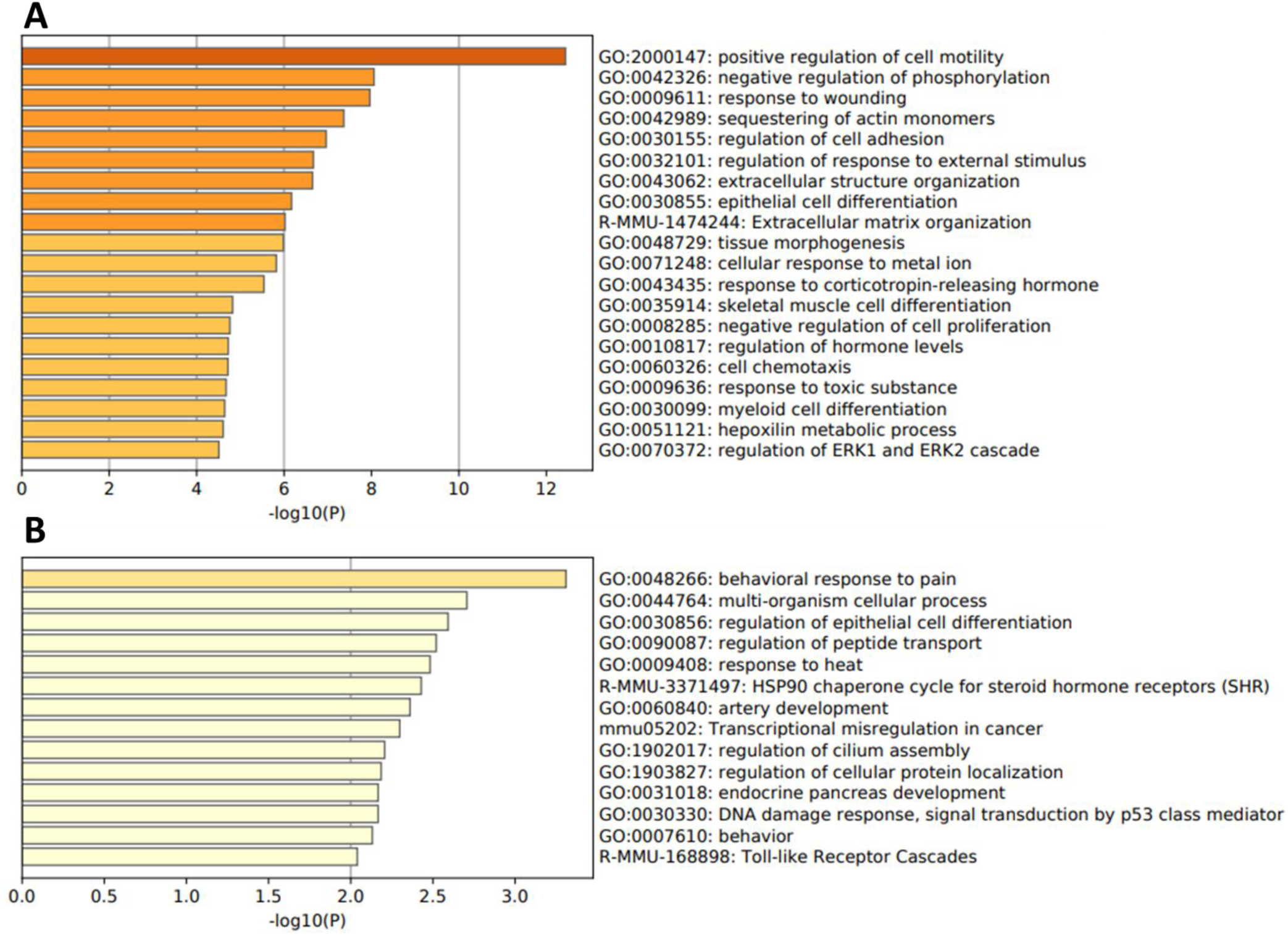
BDE-47 Gene Ontology Analysis: Top Metascape biological categories that were (A) negatively enriched by BDE-47 exposure or (B) positively enriched.

Ingenuity pathways analysis identified 77 pathways significantly enriched with -log p-values of ≥ 1.3 with all but one predicted to be suppressed (*Supplemental File 7*) (full list found at ^37^). Downregulated canonical pathways associated with pituitary endocrine function included: *GNRH signaling, TR/RXR activation, Thyroid hormone metabolism I (via Deiodination), Glucocorticoid receptor signaling, Germ cell-Sertoli cell junction signaling,* and *ERK/MAPK signaling* (*Table 3*). As with BPS, there were several altered pathways (at least 15) that were directly associated with immune function or intracellular phosphorylation cascades and downstream transcription factors (at least 35). These were typically enriched due to the downregulation of *Fos, Rasd1,* and *Gab1*.

**Table 3:** Hormone Related IPA Pathways for BDE-47 exposed pituitary. IPA canonical pathway analysis of differentially expressed genes from BDE-47 exposed mice to controls. Significantly enriched pathways determined by log p-values ≥1.3. All pathways were predicted to be downregulated.

| Ingenuity Canonical Pathways | p-value | DEGs |
| --- | --- | --- |
| GNRH Signaling | 0.0162 | *Cacna1e*, *Egr1*, *Fos*, *Gnrhr* |
| TR/RXR Activation | 0.0170 | *Dio1*, *Eno1*, *Syt12*, *Tshb* |
| ERK/MAPK Signaling | 0.0263 | *Dusp1, Elf3, Fos, Mycn, Rasd1* |
| Thyroid Hormone Metabolism I | 0.0347 | *Dio1* |
| Glucocorticoid Receptor Signaling | 0.0447 | *Dusp1, Fos, Krt17, Pgr, Rasd1* |
| Germ cell – Sertoli cell junction signaling | 0.051 | *Acta2, Actn1, Gsn, Rasd1, Tuba1c* |

The upstream regulator analysis predicted inhibition for several molecules associated with HP, including beta-estradiol (also inhibited with BPS). PgR was another predicted upstream regulator associated with the HP-gonad axis. Additionally, dexamethasone, a pharmaceutical glucocorticoid and the inflammatory cytokines, Il1B [Interleukin 1 beta], and TNF [Tumor necrosis factor], were predicted as impacted upstream regulators. Related to immune function was the predicted inhibition of the gram bacteria negative membrane component and potent pyrogen, lipopolysaccharide. Like BPS, reduction in secondary messenger of GPCR systems by BDE-47 had a significant overlap with inhibited forskolin and CREB1 activity, in addition to the important secondary messenger pathway transcription factor, Jun (*Table 4*).

**Table 4:**
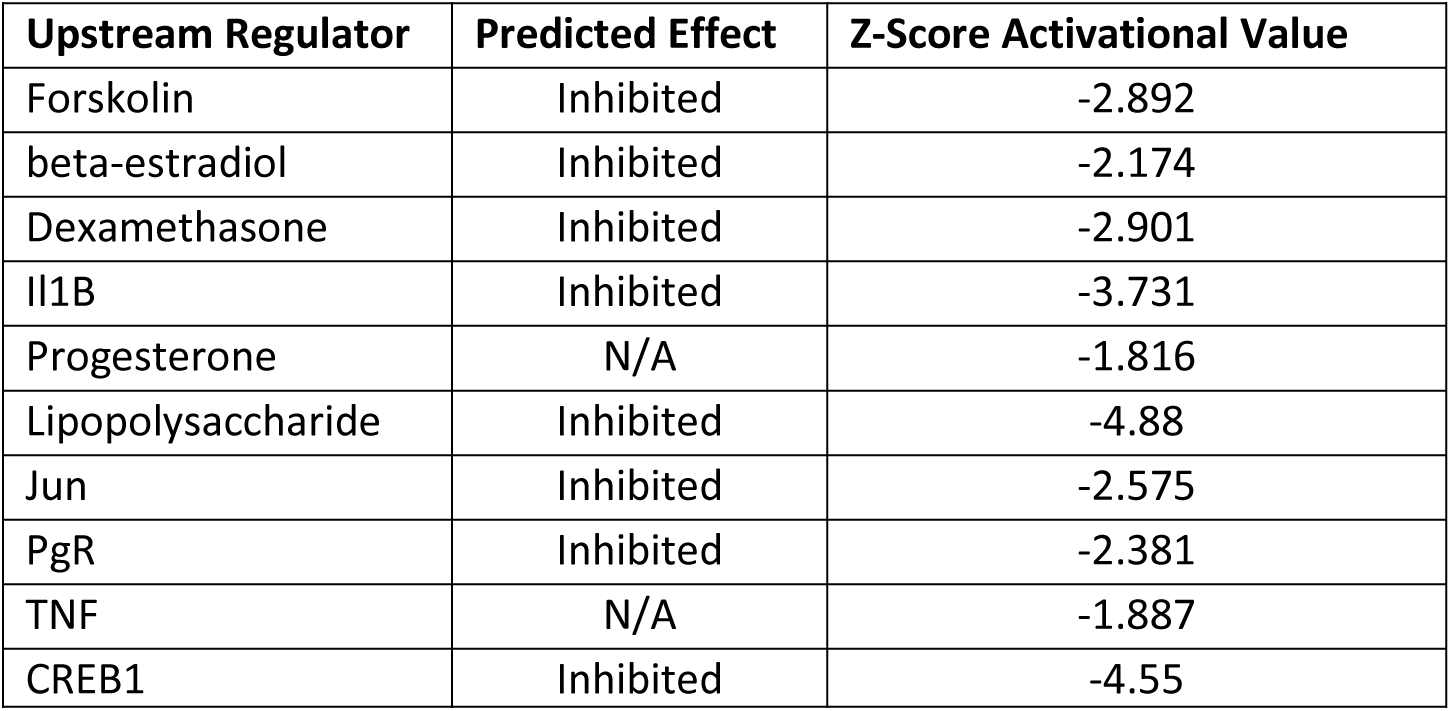
Top predicted IPA upstream regulators for BDE-47 exposed pituitaries.

| Upstream Regulator | Predicted Effect | Z-Score Activational Value |
| --- | --- | --- |
| Forskolin | Inhibited | -2.892 |
| beta-estradiol | Inhibited | -2.174 |
| Dexamethasone | Inhibited | -2.901 |
| Il1B | Inhibited | -3.731 |
| Progesterone | N/A | -1.816 |
| Lipopolysaccharide | Inhibited | -4.88 |
| Jun | Inhibited | -2.575 |
| PgR | Inhibited | -2.381 |
| TNF | N/A | -1.887 |
| CREB1 | Inhibited | -4.55 |

As with BPS, we assessed expression of key hormone regulating genes in the hypothalamus selected based on the results of IPA analysis in pituitary (*Figure 5*). None of the gonadotropin signaling related genes (*Esr1*, *Gper1, Pgrb, Kiss1*, and *Gnrh1)* were significantly altered by BDE-47. However, analysis of HP-thyroid axis signaling genes revealed that the thyroid hormone nuclear receptors, *Thra* [Thyroid hormone receptor alpha] and *Thrb* [Thyroid receptor beta], which receive and initiate negative thyroid feedback in the HP-thyroid, were both significantly downregulated. Conversely, the downstream expression of *Trh* [Thyrotropin releasing hormone], the hypothalamic inducer of pituitary thyrotropic activity, was not altered. Though IPA did not indicate that growth hormone signaling was impacted in the pituitary, previous literature demonstrated long-lasting increase in circulating insulin-like growth factor 1 (IGF-1) following perinatal exposure to BDE-47 in rats ^28^ and mice ^19^ suggesting that BDE-47 can alter the HP-liver axis. Therefore, we assessed hypothalamic genes from the growth hormone pathway. While *Sst* was unaltered, *Ghrh* was significantly upregulated in BDE-47 exposed animals. Lastly, glucocorticoid pathway target genes, *Npy* and *Crh*, were not affected by BDE-47 exposure.

**Figure 5:**
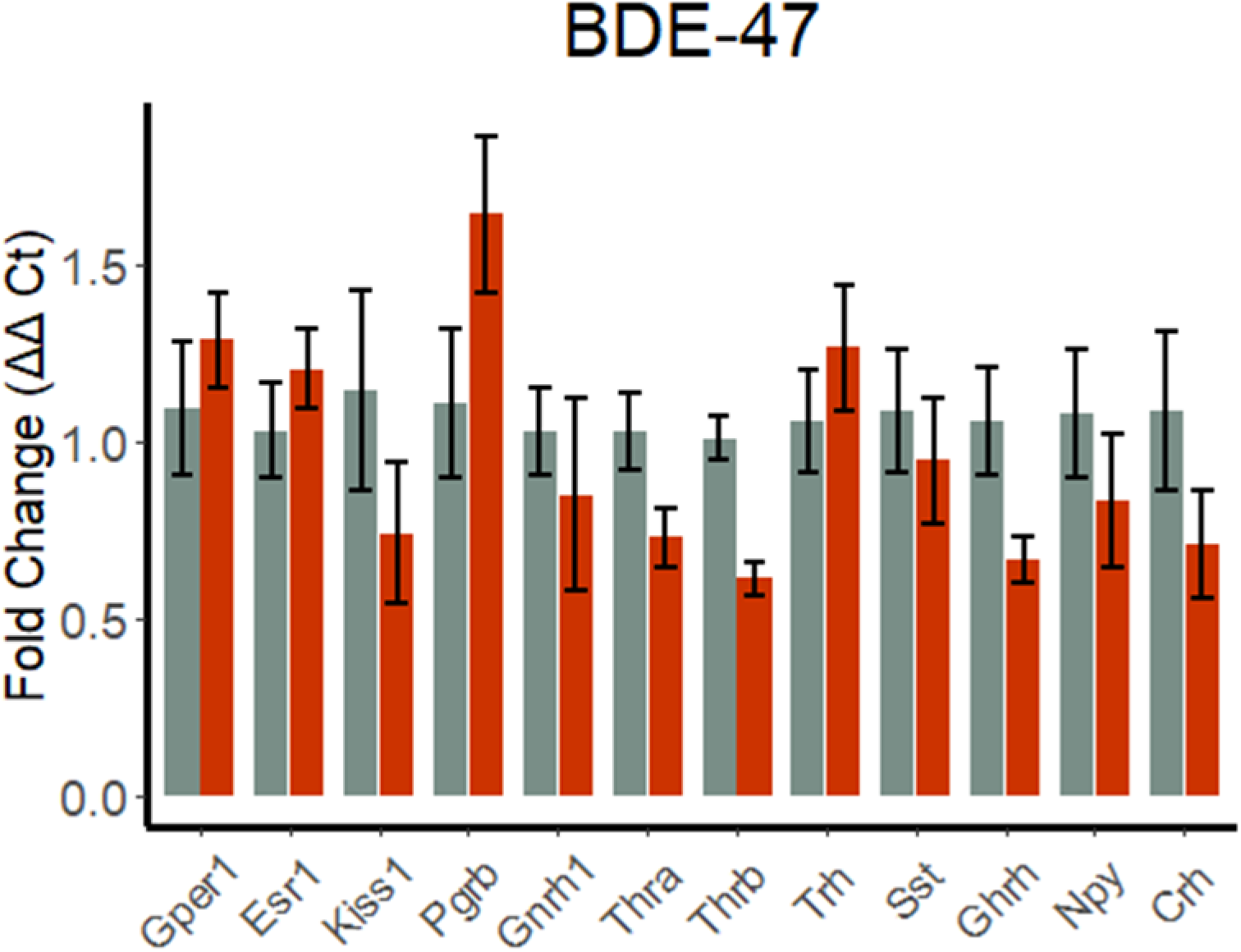
BDE-47 alters long-term hormone gene expression in the hypothalamus. Gene expression of key hormone regulating genes in hypothalamus via qPCR. Gray bars signify controls, while orange are for BDE-47 group. Significance determined by Student’s t-test or Welch’s -test where relevant, α = 0.05, * represents p < 0.05. Error bars represent standard error.

Several DEGs and altered pathways were common for BPS and BDE-57 groups, while others were unique to a group, including corticosteroid signaling in BPS group, and thyroid signaling in BDE-47 group. A summary of the major effects on hormone systems for both exposures is shown in (*Figure 6*).

**Figure 6:**
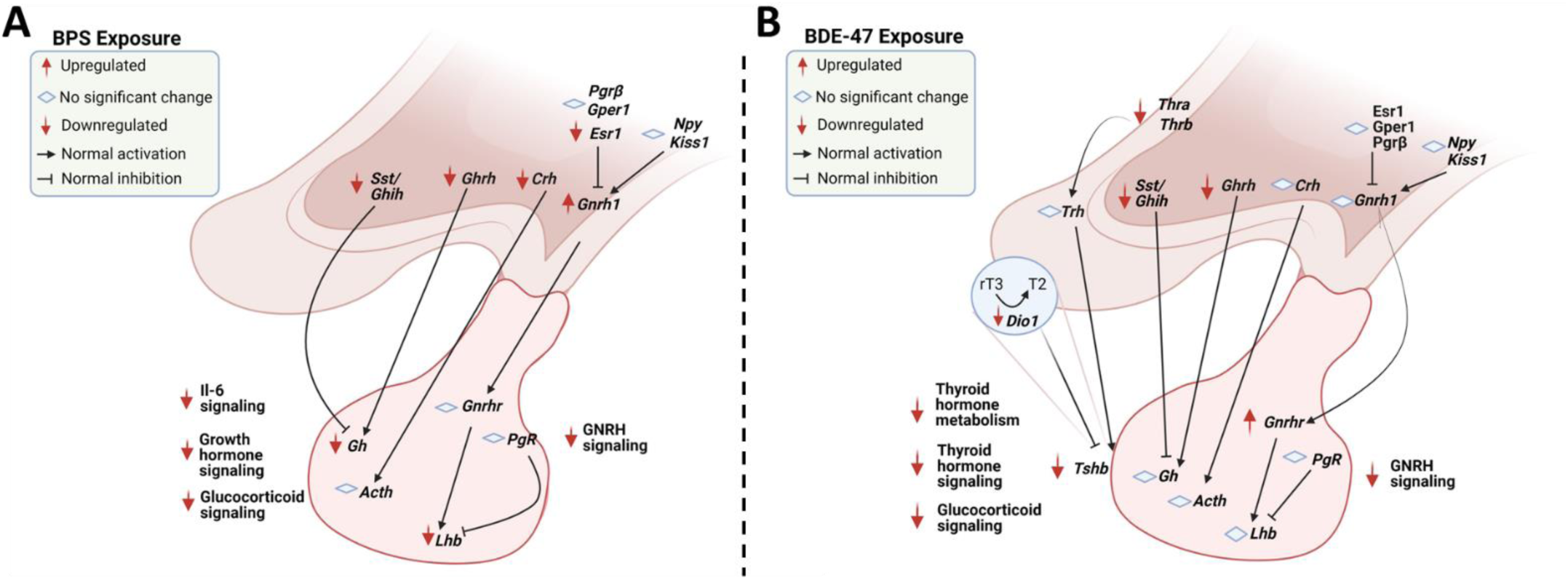
Summary of Differentially Expressed Hormone Pathways for BPS and BDE-47. Key altered hormone regulatory DEGs and pathways after exposure to BPS (A) or BDE-47 (B).

### TBBPA induces modest long-term changes in HP hormone signaling pathways

103 DEGs were identified in pituitary of TBBPA exposed mice, where 80 DEGs were significantly downregulated, and 23 DEGs were upregulated. Unlike, BPS and BDE-47, TBBPA-induced changes in gene expression affected few genes recognized as directly related to hormonal function of the pituitary. Most relevant of those effected genes, is *Igfbp2* [Insulin like growth factor binding protein 2], a circulating transport protein that binds IGFI and II proteins with high affinity and altering their activity ^46^. Also of note, the interleukin *il36G*, had downregulated expression as was seen for the previous EDCs. A full list of DEGs from the TBBPA group (*Supplemental File 8*) are found in a repository ^37^.

Metascape analysis conducted with our short list of 103 differentially expressed genes identified seven positively enriched categories (*Figure 7A*) and sixteen negatively enriched categories (*Figure 7B*). Full list found list of categories in (*Supplemental File 9*, also in repository ^37^). Notable positively enriched categories for endocrine function include *negative regulation of intracellular signal transduction* and *peptide-tyrosine phosphorylation*, which, similar to BPS and BDE-47, indicate that TBBPA alters GPCR mediated intracellular phosphorylation cascade activity. Negatively enriched pathways include at least three HP hormone related biological functions: *regulation of insulin-like growth factor (IGF) transport and uptake, neuropeptide signaling pathway*, and *retinoid metabolism and transport*.

**Figure 7:**
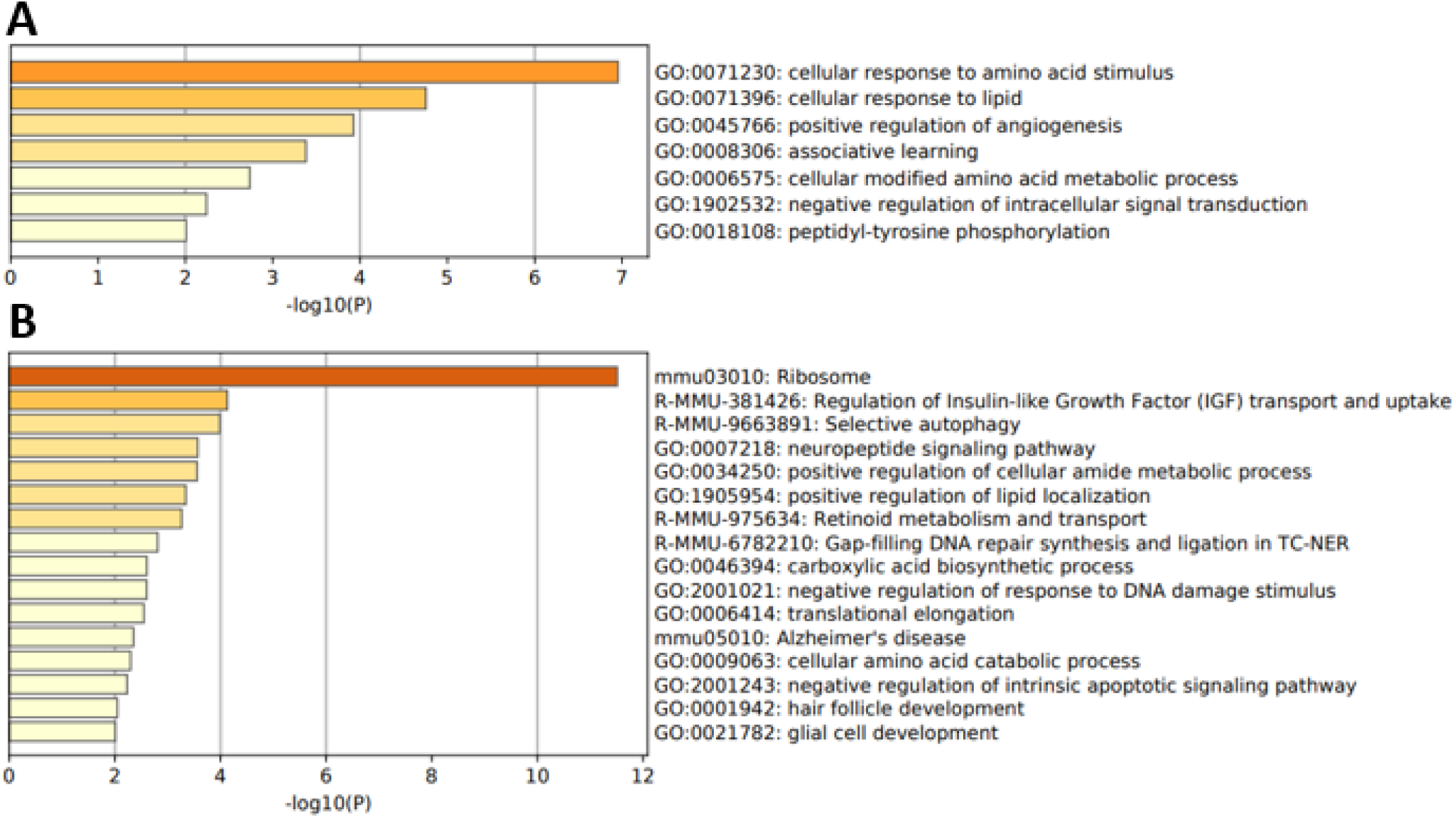
TBBPA Gene Ontology Analysis. Top Metascape Biological Categories that were (A) negatively enriched by TBBPA exposure or (B) positively enriched. Gene Ontologies were assessed from significantly differentially expressed gene list.

Ingenuity pathways analysis was conducted to evaluate predicted canonical pathway disturbances based on TBBPA exposure (*Supplemental File 10*, also in repository ^37^). All significant pathways were found to be downregulated, however, none of the pathways had obvious relevance to the hormone-signaling function of the pituitary. The most notable affected pathway was the downregulation of *EIF2* [eukaryotic initiation factor 2] signaling, which regulates eukaryotic initiation of mRNA translation. This pathway was affected by downregulation of several genes associated with 60s and 40s ribosomal subunits. Similar to BPS and BDE-47, the TBBPA group was also characterized by downregulation of immune response pathways (five total pathways) with four of these including downregulation of *il36G*.

IPA upstream analysis showed that similar to the two other EDCs, TBBPA exposure lead to predicted inhibition of the gonadal steroid, progesterone, and IGF-1 signaling, as well as inhibition of several GPCR related intracellular signaling molecules, including CREB1 and Myc signaling molecules MYCN, MYC, and MLXIPL (*Table 5*). Further supporting a reduced function of GPCR systems, the GPCR agonist, prostaglandin E2, was also inhibited. Additionally, IPA predicted inhibition of the transcription factor, TP53 [Tumor protein P53], which plays a critical role in tumor suppression and regulation of the cell cycle, and inhibition of the protein complex NFkB [Nuclear factor kappa-light-chain-enhancer of activated B cells] (complex), an important regulator of infection response, as well as the cytokine interferon gamma (IFNG).

**Table 5:** Top predicted IPA upstream regulators for TBBPA exposed pituitaries.

| Upstream Regulator | Predicted Effect | Z-score Activational Value |
| --- | --- | --- |
| MLXIPL | Inhibited | -3.564 |
| MYC | Inhibited | -3.416 |
| TP53 | Inhibited | -2.553 |
| NFkB | Inhibited | -2.449 |
| IGF1 | Inhibited | -2.399 |
| CREB1 | Inhibited | -2.392 |
| MYCN | Inhibited | -2.392 |
| IFNG | Inhibited | -2.368 |
| Progesterone | Inhibited | -2.219 |
| prostaglandin E2 | Inhibited | -2.200 |

### Convergent genetic effects of EDC exposures

Our analysis indicated that many genes, molecular pathways, and HP axes were affected by more than 1 EDC (*Figure 8*). All three exposures affected expression of 10 common genes (*Cwc22, Krt17, Vgf, Gal, Penk, Vim, Ercc2, Alox12e, IL36g/Il1f9, Junb)* with one common significantly altered pathway, acute phase signaling (due to altered expression of Il36g). BPS and BDE-47 had the greatest overlap of DEGs, 20 (*Figure 8A*), and HP-adrenal altered pathways in common (*Figure 8B*). Here, BPS and BDE-47 exposure downregulated gonadal and glucocorticoid axis pathway genes, while all three chemicals reduced growth hormone axis (HP-liver) activity. Common to these altered pathways are the ERK/MAPK signaling cascades, suggesting that these signaling molecules may have a broad sensitivity to environmental contaminants. Considering the combination of effects across our analyses in both the pituitary and hypothalamus, we summarized the key shared and specific effects of these three EDCs in (*Figure 8C*).

**Figure 8:**
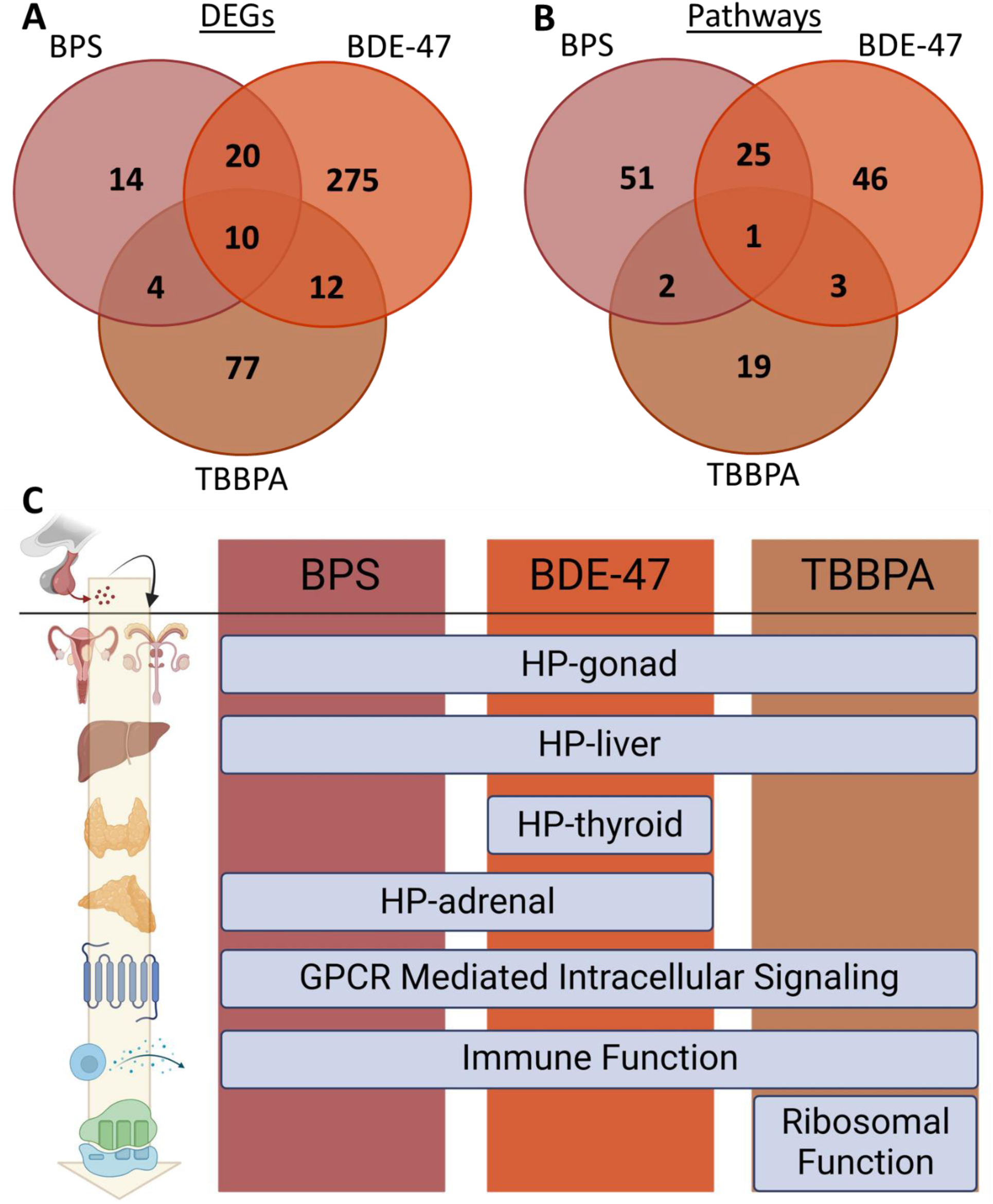
Overlap of Altered Gene Pathways due to EDC exposure. The number of shared and specific differentially expressed pituitary genes (A) and IPA altered pathways (B) between BPS, BDE-47, and TBBPA exposed individuals. (C) Summarization of EDC exposure on long-term brain hormone axis and key gene pathways. All categories were downregulated.

## Discussion

We have found that developmental exposures to three ubiquitous EDCs (BPS, BDE-47, and TBBPA) produce long term gene expression changes to central (hypothalamic-pituitary) control of endocrine functions of the organism. Many of these changes were chemical-specific, while others had partial or complete overlap across all chemical exposures. Specifically, BPS, a known estrogenic EDC with effects on downstream organs ^47,48^ showed alterations to HP-gonad gonadotropin hormone signaling. BPS also altered several other HP-related hormone pathways, such as growth hormone signaling and corticotropin signaling, that had consistently reduced expression in both the pituitary and hypothalamus. BDE-47 demonstrated suppression of the thyroid signaling axis, an effect previously seen in rodents exposed to PBDEs ^49^. Additionally, BDE-47 reduced gene expression in the gonadal and growth hormone pathways. Both results are concordant with previously reported effects of BDE-47 on reproductive outcomes ^23^ and IGF-1 in circulation ^19,28^. Figure 8 summarizes the gene and gene pathway effects of BPS (A) and BDE 47 (B). Although TBBPA exposure altered many gene pathways, particularly ribosomal proteins, there was more mild support for effects on endocrine axes, most notably a reduction in IGF binding protein regulation and steroid receptor function within the pituitary.

### Disruption of hypothalamic-pituitary-thyroid axis

BDE-47 exposure downregulated *Tshb* expression in the pituitary*. Tshb,* is the gene that encodes the glycoprotein beta subunit of thyroid-stimulating hormone (TSH), aka thyrotropin ^50^, and is responsible for inducing secretion of thyroid hormones ^51^ and HP-thyroid feedback control ^52,53^. We predicted that changes in *Tshb* expression may be related to alterations in thyroid hormone feedback genes in the hypothalamus. Indeed, we found that both isoforms of the thyroid hormone receptor, *Thra* and *Thrb*, had downregulated expression. In the pituitary, BDE-47 also downregulated *Dio1* expression, this gene codes for deiodinase type 1 ^54^, a key driver of T4 to T3 metabolism within the pituitary ^55^. Conversion of T4 to T3 is necessary in the pituitary as T3 interacts with thyroid hormone receptors to promote negative feedback of *Tshb* ^51,56^. With abnormally reduced DIO1, pituitary recognition of circulating thyroid hormone levels would be diminished. Previous studies have seen similar effects of altered DIO1 signaling with BDE-47, for example, in larval zebrafish ^57^ and rats ^58^. All together changes at hypothalamic and pituitary levels observed in this study suggest potentially weakened negative feedback control in HP-thyroid axis following developmental exposure to BDE-47.

Indeed, PBDEs and specifically BDE-47 have been well studied for their effects on thyroid disruption and as neurotoxic substances ^59–64^. The most well-documented mechanism of thyroid toxicity of PBDEs consists of thyroid hormones (THs) displacement by these substances and their metabolites from TH transport proteins: transthyretin (TTR) and thyroxine-binding globulin (TBG) ^65,66^. Based on this evidence one potential explanation of the observed weakening of the negative feedback control in HP-thyroid axis may be due to some compensatory changes during development when the axis was overstimulated due to THs displacement from transport proteins by BDE-47. Analysis of PBDE effects on thyroid signaling is complicated however, as dose-response relationships for thyroid outcomes are likely U-shaped within the range of human exposures in the general populations. For example, in a recent meta-analysis of 16 population studies, serum thyroid hormones (THs) were negatively associated with serum PBDEs when median levels of PBDEs were <30 ng/g lipid while associations were mostly positive if the median levels of PBDEs were > 100 ng/g lipids ^67^. HP-thyroid axis development is important for cognitive and neurological function and disruption to thyroid signaling may have profound effects on lifetime health ^68^. Specifically, reduction in TSH production would likely result in downregulation of thyroid hormones leading to deregulated energy and growth maintenance in the body, often exhibited in a condition called secondary hypothyroidism in humans ^69^.

TBBPA and BPS did not show any overt effects on thyroid axis gene expression. This finding is seemingly discordant to previous studies in zebrafish which have shown that TBBPA induced reduction in *Tshb* ^70^ and BPS induced increase in Deiodinase 1 and TSH-β protein ^71^ or altered thyroid function in fish ^72–74^ and rat pituitary cells ^75^. However, these studies analyzed direct effects of exposure while our experiments targeted long-lasting reprograming of HP control. This difference in the timing of exposure and outcome analysis is a likely explanation of the discrepant results between studies, although use of different models and dosing regimens may also be an important factor.

### Disruption of hypothalamic-pituitary-gonadal axis

The pituitary serves as an important mediator of gonadal feedback systems (HP-gonad) and exerts gonadal endocrine control through the release of gonadotropic hormones luteinizing hormone (LH) and follicle-stimulating hormone (FSH) ^76,77^. We found that BPS exposure downregulated expression of luteinizing hormone glycoprotein subunit β (*Lhb*). LH plays a central role as a tropic hormone that organizes sex steroid secretion in males and females ^78^. The reduction of *Lhb* and potentially LH protein, would suggest low levels of testicular steroidogenesis, deceased testosterone, and altered rate of spermiogenesis ^79–82^. An effect seen previously for the structurally similar, BPA ^83^. Though BDE-47 did not alter *Lhb* expression, it affect expression of upstream pituitary genes critical to regulation of *Lhb*. Indeed, *Gnrhr* was upregulated, a possible feedback response to decreased GNRH, while the zinc-finger transcription factor *Egr1* (early-growth repsonse protein 1) ^84^ had reduced gene expression after BDE-47 exposure, the latter of which serves as a direct mediator between *Gnrhr* and *Lhb* expression ^40^.

Reduced HP-gonad function was supported by negative enrichment of the gonadotropin-releasing hormone (GNRH) pathway for both BPS and BDE-47. Given that GNRH activation of gonadotropin-releasing hormone receptor (GNRHR) drives expression and secretion of LHβ in the pituitary ^85^, suppression of this pathway by BPS and BDE-47 was concordant with decreased *Lhb* expression in both exposed groups. In addition, IPA showed significant overlap of our HP-gonad related DEGs and the expected expression reduction seen from GNRH-A, a pharmaceutical GNRHR agonist ^86,87^. When directly assessed in our study, we found that expression of the gene for GNRH (*Gnrh1)* was altered in the hypothalamus due to BPS exposure, similar to a previous finding in rats ^88^. Conversely, BDE-47 did not have any long-term effect on hypothalamus *Gnrh1* expression. This is interesting as BDE-47 increased *Gnrhr* in the pituitary, suggesting a negative feedback response to a lack of GNRH. The lack of change in *Gnrh1* expression suggests that BDE-47 may have some direct effect on pituitary *Gnrhr* expression specifically, or by altering feedback sensitivity to gonadal steroids within only the pituitary.

To ensure brain negative feedback response to circulating gonadal hormones, the hypothalamus expresses estrogen and progesterone receptors. Given that both BPS and BDE-47 altered expression of estrogenic signaling cascade genes in pituitary, we assessed gene expression of these receptors in the hypothalamus. Indeed, gene expression of estrogen receptor isoform α (*Esr1*) was significantly downregulated, though expression of the estrogen membrane receptor (*Gper1)* and progesterone receptor (*Pgrb)* were not altered by BPS exposure. A previous study in the mammary gland indicated that estrogens can alter *Esr1* levels in the mammary and uterus ^89^, supporting our hypothesis that BPS can alter long-term gene expression through estrogenic activity, however, the response of *Gper1 to* developmental estrogens is less clear. Our data here compliments previous evidence that BPS acts as an estrogen disruptor in body endocrine glands ^47,90,91^ and other levels of the HPG axis signaling in fish ^92^, mice ^14^, and other organisms (Reviewed in Rochester, 2015 ^47^).

In the BDE-47 group no changes in expression of *Esr1, Gper1,* and *Pgrβ* were observed. Previous studies in humans have shown inconsistent effects of BDE-47 on HP-gonad endpoints, with both positive ^93^ and negative associations ^94^.

### Disruption of hypothalamic-pituitary-liver axis

HP control over secretion of growth hormone (GH), and its downstream target insulin-like growth factor 1 (IGF-1), is a central mechanisms of growth and metabolism regulation ^95,96^. Here we show that both BPS and BDE-47 downregulated expression of genes in this axis.

BPS reduced RNA expression of *Gh* and inhibited GH pathways in the pituitary, as well as reduced expression of *Ghrh* in the hypothalamus. It is also important to note that *Sst*, the gene that encodes somatostatin (aka growth hormone inhibiting hormone) was also downregulated in the pituitary by BPS suggesting reduced downstream expression of IGF-1 protein in target tissues. This is supported by the IPA finding that IGF-1 signaling was reduced in our pituitary study. Gonadal steroids are known to influence the HP-liver ^97^ and estrogens have been shown to reduce pituitary sensitivity to growth hormone-releasing hormone ^98^, suggesting that BPS effects on HPL axis may be mediated via estrogenic signaling.

BDE-47 also decreased hypothalamic *Ghrh* gene expression, however pituitary *Gh* expression was not altered by the exposure. Previous research demonstrated long term increase in IGF-1 plasma levels in rats ^28^ and mice ^19^ following developmental exposures to BDE-47. Taken together these studies indicate that developmental exposures to environmentally relevant doses of BDE-47 permanently disrupts HPL axis of hormonal regulation.

Though the *Gh* gene was not a DEG for the TBBPA exposure, IGF-1 was a predicted upstream negative regulator, suggesting that as with BPS, the HP-liver system was likely inhibited after TBBPA exposure at the pituitary level.

### Disruption of the hypothalamic-pituitary-adrenal axis

Our results demonstrated that BPS and BDE-47 inhibit the long-term activity of the HP-adrenal axis. BPS was implicated in downregulating this axis at the level of both glucocorticoid receptor and corticosteroid releasing hormone (CRH) signaling, with the former being driven principally by the secondary signaling molecules *Dusp1, Rasd1,* and *Fos.* Downregulation of glucocorticoid receptor (GR) signaling is also supported by decreased expression of *Krt17*, a cytoskeletal filament regulated downstream of GR ^99^. This was matched by a decrease in *Crh* (corticosteroid releasing hormone) in the hypothalamus, suggesting a systemic inhibition of HP-adrenal brain activity. Concordant with downregulation of *Crh,* expression of the orphan receptor and transcription factor of the nuclear receptor subfamily A1 (*Nr4A1*) was significantly decreased in the pituitary. Nr4A1 has an important role in mediating HP-adrenal negative feedback and has been used previously as a marker of neuronal stress ^45^. In response to stress, Nr4A1 expression in the hypothalamus is increased by CRH in the hypothalamus as well as the pituitary. Conversely, circulating adrenal corticosteroids negatively regulate Nr4A1, either along with CRH or in result of lower CRH ^45^. Our findings are consistent with elevated circulating corticosteroid in the serum brought on by the early BPS exposure, as seen previously in adolescent rats following developmental exposure to the bisphenol BPA ^100^.

Pathway analysis indicated that pituitary HP-adrenal feedback control was generally inhibited by BDE-47 exposure. Specifically, BDE-47 downregulated *Krt17* and progesterone receptor (PgR). PgR was shown before to regulate stress response ^101^. Indeed, humans provided CRH or ACTH, have significant increases in plasma progesterone levels ^102^, supporting our findings that BDE-47 exposure alters both corticosteroid and progesterone signaling. Elevated progesterone secretion from the adrenals ^103^ likely acts upon PgR in the brain to regulate feedback on CRH expression, as suggested by a study in placental primary culture ^104^. The effect that progesterone has on the HP-adrenal axis is best studied in the rodent brain where progesterone’s metabolite, allopregnanolone, modulates GABA-A receptors to reduce stress-induced anxiety ^101^. However, a study in rats indicated progesterone’s direct interaction with PgR reduced anxious behavior suggesting that the receptor has a direct role in reducing HP-adrenal stress ^105^. BDE-47’s inhibitory effect on *Pgr* expression, a typical consequence of circulating corticosteroids, corresponds to inhibition of CRH and GR signaling seen in the BPS group and suggests the likelihood of long-term glucocorticoid blood elevation by both EDCs.

### GPCR-mediated phosphorylation cascades and immune system are novel targets for convergent endocrine disruptors

Several intracellular signaling molecules downstream of G-protein coupled receptors (GPCRs), including phosphorylating molecules and transcription factors, were affected by both BPS and BDE-47 exposures. For example, *Rasd1* ^106^ and *Dusp1* ^107^, signal transducers of GPCRs and MAPK signaling cascades, were downregulated in both BDE-47 and BPS exposed groups. Furthermore, both exposures also downregulated the transcription factors, *Fos* and *Junb* expression, which form AP-1 complexes ^108^ that are important for driving transcription for many cellular receptor systems, including nuclear receptors and GPCRs like GNRHR ^109^ and GPER ^110^. One GPER, GPER1 (G protein-coupled estrogen receptor 1), an estrogen receptor that stimulates adenylate cyclase and cyclic AMP, was predicted to be an altered upstream regulator in BPS exposed mice. However, GPER1 was not directly altered in either the pituitary or hypothalamus. *Egr-1* expression, which codes for a zinc-finger transcription factor responsive to environmental stressors and cancer ^111^, was also downregulated by BDE-47 and BPS. TBBPA exposure did not result in the same altered signal transduction profile, but it did reduce expression of one major transcription factor, *Myc*, and had IPA predicted inhibition of the GPCR intracellular signaling molecules MYC, TP53, CREB1, MYCN as well as the GPCR agonist prostaglandin E2.

The altered expression of genes shown here code for proteins that serve intracellular signaling cascades that can be triggered by multiple membrane receptor ligands and intermembrane signaling machinery^112^. In fact, BPS, BDE-47, and TBBPA had overlapping effects on predicted pathways primarily through mRNA alteration of intracellular signaling molecules and downstream transcription factors that are ubiquitously expressed in tissues. Here we see that these generalized intracellular molecules serve as common templates for which environmental chemicals can alter gene transcription, even if biochemical class and structure is different ^9^. These factors may serve as a point of wide disrupting convergence for many chemicals and deserves further attention. An unexpected finding of this study was the consistent alteration, specifically reduction, in immune signaling functions in the pituitary by all three chemicals. Notably, BPS and BDE-47 shared inhibitory effects on tumor necrosis factor (TNF) while all three EDCs had predicted inhibition of nuclear factor kappa-light-chain-enhancer of activated B cells (NF-kB) signaling, which are important pathways for immune response ^113^. Innate immune responses such as Toll-like receptor signaling were downregulated by BPS and BDE-47. Shared DEGs of these exposures included reduced *Fos* and *Il36G* (codes for interleukin 36 G), while BPS had additional reduction of the toll-like receptor family gene, *Tlr1*. TBBPA also had Il36G as a negatively regulated DEG but lacked alterations to FOS or other DEGs to give significant change in the IPA Toll-like Receptor signaling pathway. However, TBBPA did alter several other immune pathways (available in repository ^37^). It is interesting that TBBPA had reduced immune molecule signaling, similar to BPS and BDE-47, but without indication of change in corticosteroid signaling as the other two did. This may suggest that immune signaling response is more generally sensitive to EDCs’ exposures, and can occur in the absence of altered corticosteroid signaling.

Altogether, the convergent disruption to both innate and adaptive immune systems in the pituitary, from multiple distinct chemicals, offers an interesting foundation for future study. Though congealing of a recognized “hypothalamic pituitary immune axis” has not yet occurred within the field of endocrinology, the interplay of both gonadal and thyroidal hormones, as well as chemical disruptors of these, has been recognized to alter immune function ^114,115^. These results warrant further investigation into both short and long-term immune responses in the pituitary under contaminant exposure.

## Conclusion

The ubiquity and variety of endocrine disrupting chemicals in the environment poses a wide potential for risk to hormone regulation of development and homeostasis. EDCs have been well studied for effects on single organ targets, but little is known about chemicals’ effects on hierarchical control of hormone regulation at the HP level. Here we show that developmental exposures to three structurally different EDCs: BPS, BDE-47, and TBBPA, can permanently alter pituitary expression of genes responsible for regulating all endocrine axes. Importantly, some hormone axes, such as the HP-gonad and HP-liver, had inhibited activity across all three chemicals suggesting a special sensitivity of these systems to chemical exposures and priority targets for future EDC study. Furthermore, these convergent chemical effects may be explained, at least in part, by the consistent reduction in GPCR mediated intracellular secondary signaling cascades seen across each EDC exposure. Specifically, the activities of RASD1, DUSP1, MYC, FOS, JUN, and CREB1 were consistently downregulated across exposures. Therefore, these molecules and other members of their signaling pathways may prove to be common targets, and important biomarkers, for EDC exposures generally. Additionally, this study has also provided novel evidence that both innate and adaptive immune cell signaling molecules in the pituitary (especially Il36G) are shared, sensitive, novel targets of EDCs’ exposure. Ultimately, this study provides evidence of shared endocrine disruption of the hypothalamic-pituitary axis system and identifies common pathways affected by EDCs.

