## Supplementary figures and images for "Developmental Reprogramming of Hypothalamic-Pituitary Axis in Mice by Common Environmental Pollutants"

### Supplemental File 1

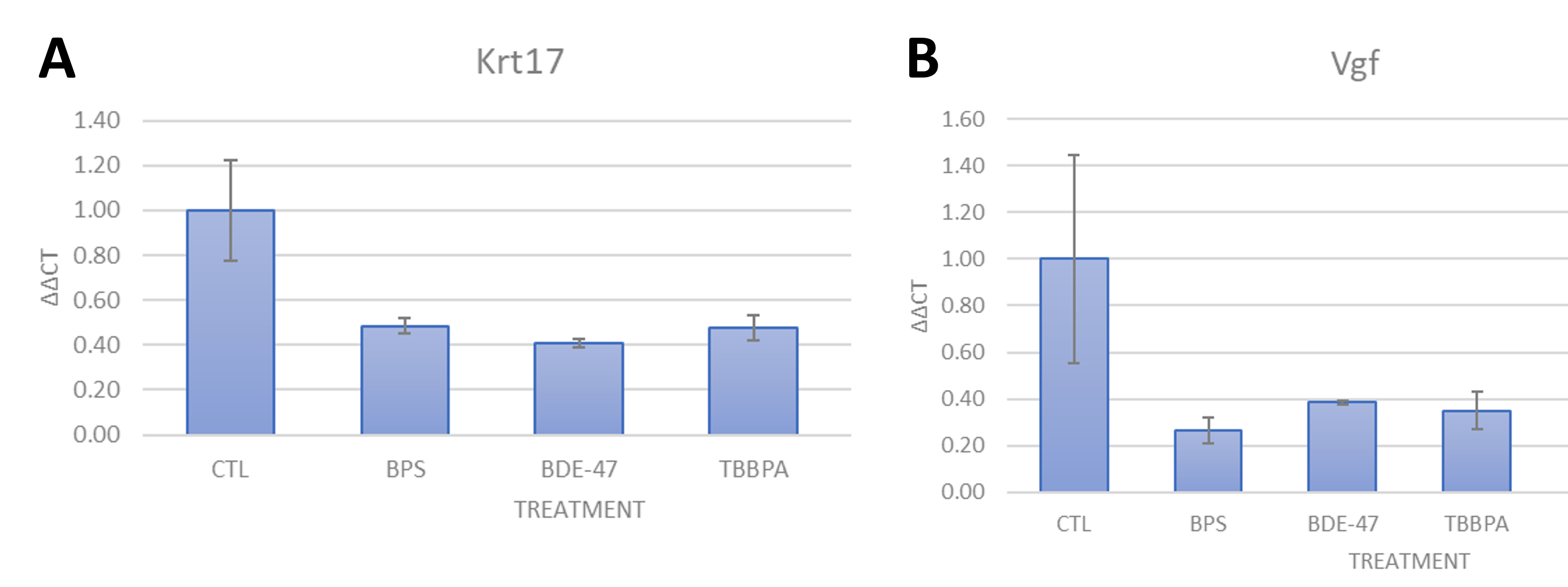
